# Atypical TGF-β Signaling Controls Neuronal Guidance in *Caenorhabditis elegans*

**DOI:** 10.1101/2020.09.15.297556

**Authors:** Oguzhan Baltaci, Mikael Egebjerg Pedersen, Tessa Sherry, Ava Handley, Goda Snieckute, Matilda Haas, Stuart Archer, Roger Pocock

## Abstract

Coordinated expression of cell adhesion and signaling molecules is crucial for brain development. Here, we report that the *Caenorhabditis elegans* transforming growth factor-beta (TGF-β) type I receptor SMA-6 (small-6) acts independently of its cognate TGF-β type II receptor DAF-4 (dauer formation-defective-4) to control neuronal guidance. SMA-6 directs neuronal development from the epidermis through interactions with three, orphan, TGF-β ligands. Intracellular signaling downstream of SMA-6 limits expression of NLR-1, an essential Neurexin-like cell adhesion receptor, to enable neuronal guidance. Together, our data identify an atypical TGF-β-mediated regulatory mechanism to ensure correct development of the nervous system.

## Main Text

Brain development requires the generation and assembly of neurons into interconnected circuits. During these highly choreographed processes, neurons are guided by interactions with adjacent cells and the extracellular matrix, which present a variety of cell adhesion and signaling molecules. Among these are the transforming growth factor beta (TGF-β) family of cytokines that are critical for metazoan development (*1*). TGF-β ligands control cell signaling by binding to and coordinating cell surface dimerization of type I and type II TGF-β receptor serine/threonine kinases (*2*). Upon dimerization, the type II receptor phosphorylates the type I receptor kinase, which transmits the signal by phosphorylating cytoplasmic SMAD (small and mothers against decapentaplegic) proteins (*2*). Once phosphorylated, SMAD proteins form complexes that translocate to the nucleus where they interact with nuclear cofactors to regulate target gene transcription (*2*).

*Caenorhabditis elegans* encodes a single TGF-β type II receptor (DAF-4) that acts with two TGF-β type I receptors: SMA-6 to control body size/male tail development, and DAF-1 to regulate dauer formation (Fig. 1A) (*3, 4*). The TGF-β ligands DBL-1 and DAF-7 control the SMA-6 and DAF-1 signaling pathways, respectively (Fig. 1A) (*5, 6*). However, the function for other putative TGF-β ligands, TIG-2 and TIG-3, are undescribed and UNC-129 controls dorso-ventral axon guidance independently of TGF-β receptors (Fig. 1A) (*7*). Transcriptional output from each *C. elegans* TGF-β pathway is regulated by specific SMAD proteins: SMA-2/3/4 (SMA-6 signaling) and DAF-3/8/14 (DAF-1 signaling) (*8*).

**Figure 1.**
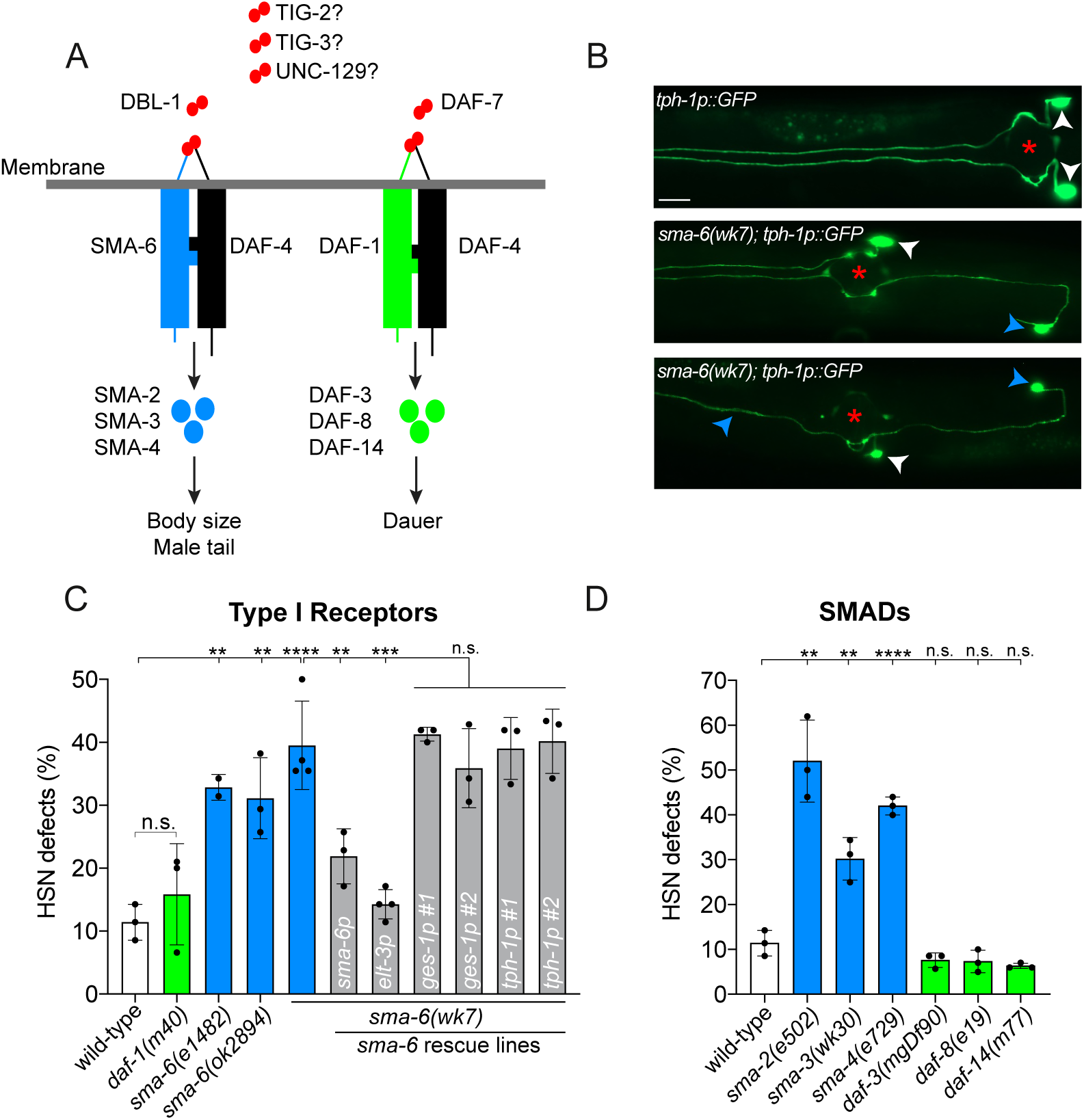
SMA-6, a TGF-β type I receptor controls HSN development. (A) The TGF-β signaling pathways in *C. elegans* control body size and male tail development (left), and dauer formation (right). Each pathway utilizes a common TGF-β type II receptor (DAF-4), and distinct ligands (DBL-1 and DAF-7), TGF-β type I receptors (SMA-6 and DAF-1) and SMAD transcriptional regulators (SMA-2/3/4 and DAF-3/8/14). TIG-2, TIG-3 and UNC-129 are orphan ligands that have not been assigned to either pathway. Receptors shown as monomers for simplicity. (B) HSN anatomy of wild-type and *sma-6(wk7)* mutant animals (fluorescent micrographs of *zdIs13(tph-1p::GFP)* labelled HSNs). In wild-type animals (top panel), HSN cell bodies migrate just posterior to the vulva and extend axons into separate fascicles in the ventral nerve cord. In *sma-6(wk7)* animals (middle and bottom panels), HSN cell bodies do not migrate correctly and their axons are misguided. Vulval position is marked with a red asterisk, wild-type positioned cell bodies with white arrowheads, and misguided cell bodies and axons with blue arrowheads. Ventral view, anterior to the left. Scale bar: 20 μm. (C) Quantification of HSN developmental defects in TGF-β type I receptor mutants *sma-6* and *daf-1*. Loss of *sma-6* but not *daf-1* causes HSN developmental defects. Driving *sma-6* expression using its own promoter or a heterologous hypodermal promoter (*elt-3*) rescues *sma-6(wk7)*-induced HSN developmental defects. Driving *sma-6* expression using intestinal (*ges-1*) or HSN (*tph-1*) promoters does not rescue *sma-6(wk7)*-induced HSN developmental defects. # refers to independent transgenic lines. n>100; **P<0.01, ***P<0.001, ****P<0.0001, n.s. not significant (One-way ANOVA with Tukey’s correction). Error bars represent mean ± SEM. (D) Quantification of HSN developmental defects in animals lacking SMAD transcriptional regulators. Loss of SMADs that control body size and male tail development (SMA-2/3/4) but not dauer SMADs (DAF-3/8/14) causes HSN developmental defects. n>100; **P<0.01, ****P<0.0001, n.s. not significant (One-way ANOVA with Tukey’s correction). Error bars represent mean ± SEM.

How TGF-β signaling coordinates gene expression programs to control nervous system development is not fully understood. Here, we used the *C. elegans* hermaphrodite-specific neurons (HSNs) to explore the *in vivo* requirement for TGF-β signalling in neuronal migration and axon guidance. During embryogenesis, the HSNs undergo long-range migration through a field of epidermal cells. Later, in juvenile larvae the HSNs extend axons over a precisely defined route within the ventral nerve cord before terminating at the nerve ring (Fig. 1B) (*9, 10*). These developmental events are precisely regulated by multiple conserved guidance pathways and environmental factors (*10-14*). We examined HSN development using a transgenic strain in which the HSN cell bodies and axons are marked with green fluorescent protein (GFP) (Fig. 1B). We tested the effect of independently disrupting the two recognized TGF-β pathways in *C. elegans* using mutants for the TGF-β type I receptors SMA-6 (body size/male tail development pathway) and DAF-1 (dauer pathway) (Fig. 1A-C). We found that SMA-6, but not DAF-1, is required for HSN migration and axon guidance (Fig. 1B-C and Table S1). Analysis of three independently-derived *sma-6* loss-of-function alleles confirmed the importance of SMA-6 for HSN development (Fig. 1C). SMA-6 is primarily expressed in the hypodermis (epidermis), from which it regulates body size, and the intestine (*15, 16*). To examine where SMA-6 acts to control HSN development, we restored *sma-6* expression under its own promoter and tissue specific drivers in *sma-6(wk7*) mutant animals (Fig. 1C). Expression of *sma-6* cDNA under its own promoter or a hypodermal promoter (*elt-3*) rescued the *sma-6(wk7*) HSN developmental defects (Fig. 1C and Table S1) (*17*). Expression of *sma-6* in the intestine (*ges-1* promoter) or HSNs (*tph-1* promoter) did not rescue HSN developmental defects (Fig. 1C and Table S1) (*18, 19*). These data show that SMA-6 functions in the hypodermis to non-cell-autonomously control HSN development.

TGF-β type I receptors control the phosphorylation status of downstream SMAD transcriptional regulators. Phosphorylation of SMADs permit their nuclear accumulation where they activate or repress target gene expression. SMADs acting downstream of SMA-6 to control body size and male tail development are SMA-2, SMA-3 and SMA-4 (*8*). We found that loss of *sma-2, sma-3* or *sma-4* caused similar penetrance of HSN developmental defects to *sma-6* mutant animals (Fig. 1C-D and Table S1). In contrast, SMADs that act downstream of DAF-1 in the dauer pathway (DAF-3, DAF-8 and DAF-14) are dispensable for HSN development (Fig. 1D) (*8*). These data show that SMA-6-directed regulation of HSN development uses the canonical transcription factors SMA-2/3/4.

The SMA-6 TGF-β type I receptor functions in a complex with the sole DAF-4 TGF-β type II receptor to control body size and male tail morphogenesis (*8*). Surprisingly, two widely used *daf-4* deletion alleles - *e1364* and *ok828* - exhibited wild-type HSN development (Fig. 2A-B). These *daf-4* alleles harbor large deletions that lead to premature stop codons and, in the case of *e1364*, elimination of kinase function (Fig. 2A and Fig. S1) (*20*). To confirm that DAF-4 is dispensable for HSN development, we generated two additional deletion alleles using CRISPR-Cas9 that introduce frameshifts in exon 1 of *daf-4* (*rp122* and *rp123*) (Fig. 2A and Fig. S1) (*21*). HSNs developed normally in *rp122* and *rp123* animals confirming that DAF-4 does not control HSN development (Fig. 2B).

**Figure 2.**
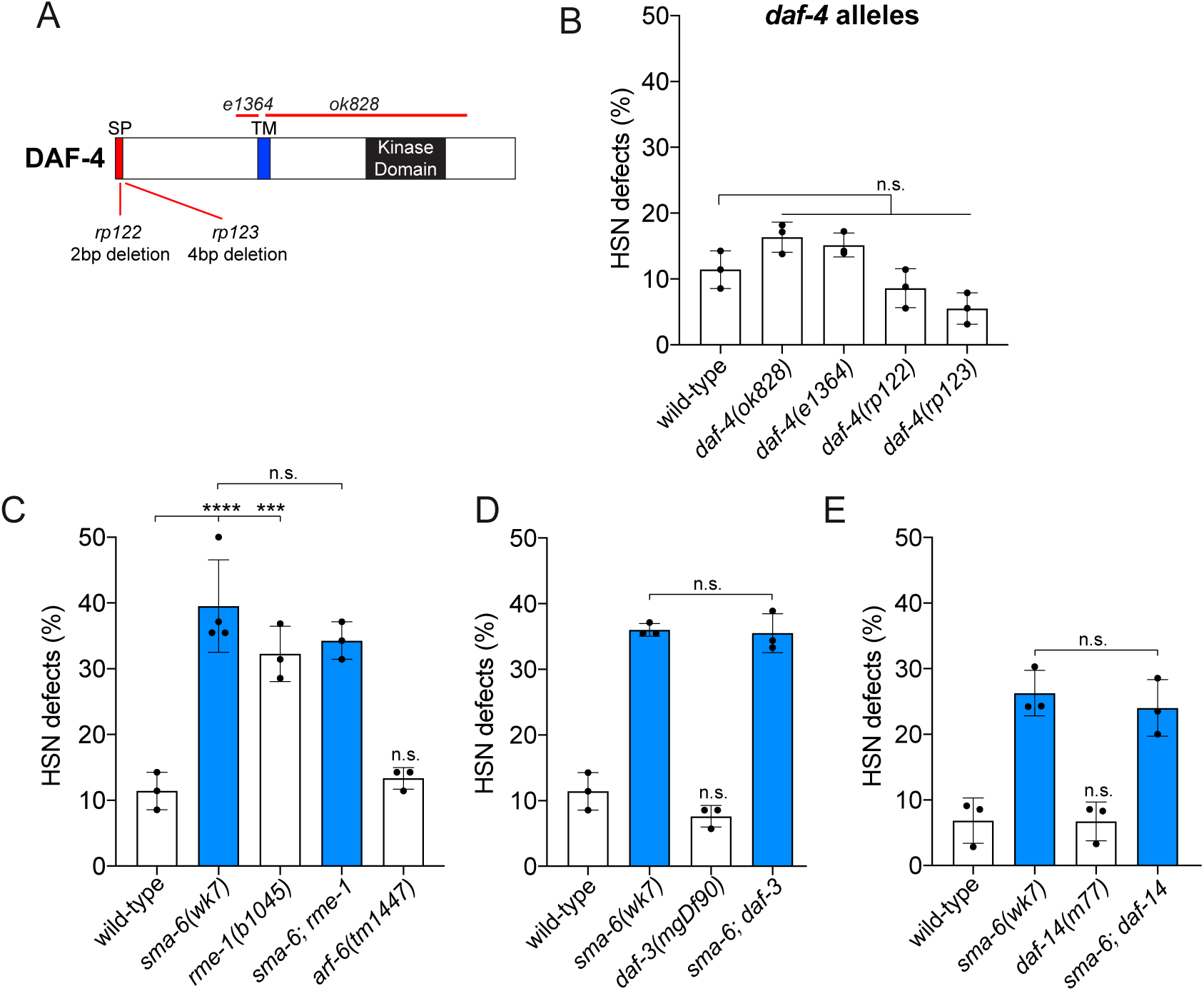
DAF-4, the sole *C. elegans* TGF-β type II receptor, is dispensable for HSN development. (A) Protein domain organisation of the DAF-4 TGF-β type II receptor and mutant alleles used in this study. All genetic lesions (red lines) cause frameshifts. SP = signal peptide; TM = transmembrane domain. (B) Quantification of HSN developmental defects in wild-type and *daf-4* mutant animals. All *daf-4* mutant alleles exhibit wild-type HSN development. n>100; n.s. not significant (One-way ANOVA with Tukey’s correction). Error bars represent mean ± SEM. (C) Quantification of HSN developmental defects in wild-type, *sma-6(wk7), rme-1(b1045), arf-6(tm1447)* and *rme-1; sma-6* mutant animals. Loss of retromer-dependent SMA-6 recycling but not ARF-6-dependent DAF-4 recycling causes HSN developmental defects. n>100; ***P<0.001, ****P<0.0001, n.s. not significant (One-way ANOVA with Tukey’s correction). Error bars represent mean ± SEM. (D) Quantification of HSN developmental defects in wild-type, *sma-6(wk7), daf-3(mgDf90)* and *sma-6; daf-3* mutant animals. n>100; n.s. not significant (One-way ANOVA with Tukey’s correction). Error bars represent mean ± SEM. (E) Quantification of HSN developmental defects in wild-type, *sma-6(wk7), daf-14(m77)* and *sma-6; daf-14* mutant animals. n>100; n.s. not significant (One-way ANOVA with Tukey’s correction). Error bars represent mean ± SEM. Note: the *mgIs71(tph-1p::GFP)* transgene was used for HSN analysis in (E) which has a lower background HSN phenotype compared to *zdIs13(tph-1p::GFP)*.

Although DAF-4 is dispensable for HSN development, it is conceivable that lack of SMA-6 causes aberrant DAF-4 receptor trafficking or dysregulation of DAF-4-directed signaling. To investigate these possibilities, we first examined the effect of disrupting TGF-β receptor recycling on HSN development. Following endocytosis, TGF-β receptors are trafficked to early endosomes, from where they are either recycled to the plasma membrane for further signaling, or conveyed to the lysosome for degradation. A previous study showed that SMA-6 and DAF-4 are recycled through separate mechanisms (*17*). RME-1 (Eps15 homology-domain containing/receptor mediated endocytosis-1) is a conserved protein required for trafficking of SMA-6 and DAF-4 (*17*). Loss of *rme-1* reduces SMA-6 levels due to inappropriate lysosomal degradation and causes accumulation of DAF-4 in intracellular vesicles (*17*). In contrast, ARF-6 (ADP-ribosylation factor-6) specifically controls DAF-4 recycling, where DAF-4 incorrectly accumulates in endosomes of *arf-6* mutants (*17*). We found that *rme-1(b1045)* mutants exhibit ∼30% penetrant HSN defects, similar to that caused by loss of SMA-6 (Fig. 2C). Additionally, loss of *rme-1* did not enhance *sma-6(wk7)* HSN defects, suggesting they act in the same genetic pathway (Fig. 2C). In contrast, *arf-6(tm1447)* mutants exhibited wild-type HSN development (Fig. 2C). Next, we examined whether the HSN developmental defects of *sma-6(wk7)* animals are caused by inappropriate DAF-4 signaling. To this end, we introduced mutants for transcriptional regulators acting downstream of DAF-4 in the dauer pathway into the *sma-6(wk7)* background. We found that neither loss of the antagonistic co-SMAD DAF-3 nor the R-SMAD DAF-14 agonist altered the *sma-6(wk7)* mutant HSN developmental defects (Fig. 2D-E). Collectively, our genetic data reveal that the SMA-6 TGF-β type I receptor controls HSN development independently of signaling directed by the TGF-β type II receptor DAF-4.

Five putative TGF-β ligands (DAF-7, DBL-1, TIG-2, TIG-3 and UNC-129) are encoded by the *C. elegans* genome, with functions identified for only two of them (DAF-7 and DBL-1) in controlling TGF-β signaling (*5, 6*). We assessed which TGF-β ligand(s) control HSN development. The DAF-7 ligand acts through the DAF-1/DAF-4 receptor complex and was not required for HSN development (Fig. 1A and 3A). DBL-1 acts in the body size/male tail development pathway through the SMA-6/DAF-4 receptor complex, however, *dbl-1* was also dispensable for HSN development (Fig. 1A and 3A). We next examined the other less studied TGF-β ligands that have not previously been associated, either functionally or physically, with a TGF-β receptor. Surprisingly, we found that knockout of any one of TIG-2, TIG-3 or UNC-129 caused HSN developmental defects (Fig. 3A). Using compound mutant analysis, we examined whether these ligands act in the same genetic pathway to control HSN development. We found that all double mutant combinations and the *tig-2; tig-3; unc-129* triple mutant exhibit the same penetrance and expressivity of HSN developmental defects as the respective single mutants - suggesting they act in concert to control HSN development (Fig. 3A and Table S1). *tig-2, tig-3* and *unc-129* are prominently expressed in body wall muscle (BWM) and the nervous system (*22*). We examined whether the source of these secreted TGF-β ligands is important for HSN development. We performed tissue-specific transgenic rescue experiments by expressing cDNAs of each ligand in BWM or the nervous system in the respective mutant backgrounds (Fig. S2) (*23, 24*). We found that expressing TIG-3 and UNC-129 in BWM rescued HSN developmental defects of the respective mutant (Fig. S2). In contrast, TIG-2 expression in the nervous system rescued HSN developmental defects in *tig-2* mutant animals (Fig. S2). Further, overexpression of UNC-129 in the nervous system exacerbated HSN developmental defects of the *unc-129* mutant (Fig. S2). These data show that correct spatial expression of TGF-β ligands is important for HSN development. We performed further genetic analysis to determine whether *tig-2, tig-3* and *unc-129* act in the same genetic pathway as *sma-6* to control HSN development (Fig. 3B and Table S1). We found that the HSN developmental defects of each compound mutant between SMA-6 and the individual ligands was not significantly different from each single mutant (Fig. 3B).

**Figure 3.**
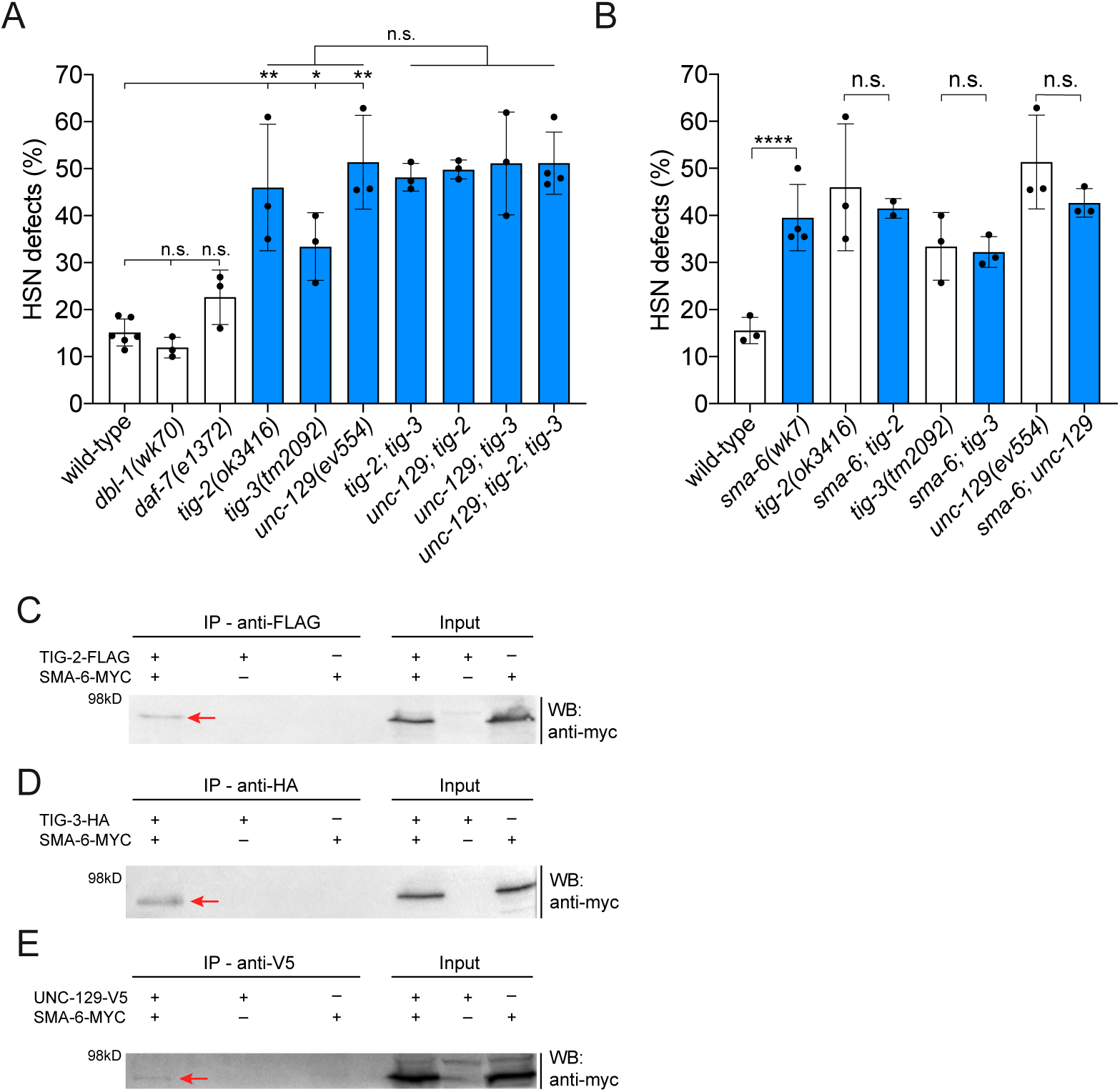
SMA-6 interacts with three, orphan, TGF-β ligands to control HSN development. (A) Quantification of HSN developmental defects in *dbl-1(wk70), daf-7(e1372), tig-2(ok3416), tig-3(tm2092)* and *unc-129(ev554)* single mutants, and all compound mutant combinations of *tig-2, tig-3* and *unc-129. dbl-1* and *daf-7* mutant animals exhibit wild-type HSN development. Loss of either *tig-2, tig-3* or *unc-129* causes HSN developmental defects and the *tig-2; tig-3; unc-129* triple mutant and each double mutant combination is not significantly different from each single mutant. n>100; *P<0.05, **P<0.01, n.s. not significant (One-way ANOVA with Tukey’s correction). Error bars represent mean ± SEM. (B) Quantification of HSN developmental defects in *sma-6(wk7)* animals in combination with either *tig-2(ok3416), tig-3(tm2092)* or *unc-129(ev554)* mutations. Each double mutant combination is not significantly different from the respective single mutant. n>100; ****P<0.0001, n.s. not significant (One-way ANOVA with Tukey’s correction). Error bars represent mean ± SEM. (C-E) Input (total cell lysates) and immunoprecipitates (IP) from transfected HEK293T cells. Proteins detected with antibodies in western blots (WB) as indicated. (C) TIG-2-FLAG co-precipitates with SMA-6-MYC; (D) TIG-3-HA co-precipitates with SMA-6-MYC; (E) UNC-129-V5 co-precipitates with SMA-6-MYC. kD, kilodalton. Immunoprecipitated SMA-6-MYC is marked with a red arrow. Whole blots in Figure. S3.

Our genetic data suggest that the TIG-2, TIG-3 and UNC-129 ligands control HSN development by interacting with the SMA-6 TGF-β type I receptor. To test this assertion, we investigated whether SMA-6 interacts with TIG-2, TIG-3 and UNC-129 using co-immunoprecipitation (coIP) experiments in human HEK293T cells (Fig. 3C-E). We found that MYC-tagged SMA-6 co-precipitated with TIG-2-FLAG, TIG-3-HA and UNC-129-V5 (Fig. 3C-E and Fig. S3). Together, our genetic and biochemical analysis show that the TGF-β ligands TIG-2, TIG-3 and UNC-129 interact with the SMA-6 TGF-β type I receptor and act in the same genetic pathway to control HSN development.

How does SMA-6-directed signaling control HSN development? To answer this question, we focused our analysis on SMA-3, an R-SMAD, that controls transcriptional outputs downstream of SMA-6 signaling and is important for HSN guidance (Fig. 1D). We performed RNA sequencing (RNA-seq) to identify differentially expressed genes (DEGs) in *sma-3(wk30)* null mutant animals compared to wild-type at the L2 stage of development, a stage during which the HSN axons extend (*9*). We identified 406 (FDR < 0.05, absolute log2Fc > 0.585) and 2082 (FDR < 0.05) DEGs in *sma-3(wk30)* animals compared to wild-type (Fig. 4A, Fig. S4-5 and Tables S2-3). Gene ontology (GO) analysis of DEGs revealed enrichment of several biological processes, including those previously associated with the regulation of body size and male tail development (Fig. S4) (*8, 25*). However, regulation of neuronal development was not an enriched GO term, suggesting that SMA-3 controls a specific gene to control HSN development. We manually surveyed the DEG lists for molecules with putative or known roles in cell signaling and adhesion - likely architects of HSN development. We identified eleven such DEGs, two downregulated and nine upregulated in *sma-3(wk30)* animals (Fig. 4A, Figs. S4-5 and Table S4). To assess functional importance of these DEGs in HSN development, we used RNA-mediated interference (RNAi) (Fig. 4A-C). For the genes downregulated in *sma-3(wk30)* animals, *sgk-1* and *mboa-6*, we knocked down their expression in wild-type animals but no HSN developmental defects were detected (Fig. 4B). To assess the potential impact of inappropriate expression of the nine genes upregulated in *sma-3(wk30)* animals, we knocked down their expression in the *sma-3(wk30)* mutant and asked whether HSN developmental defects were suppressed (Fig. 4C). We found that RNAi knockdown of *nlr-1* suppressed the HSN defects of *sma-3(wk30)* animals (Fig. 4C).

**Figure 4.**
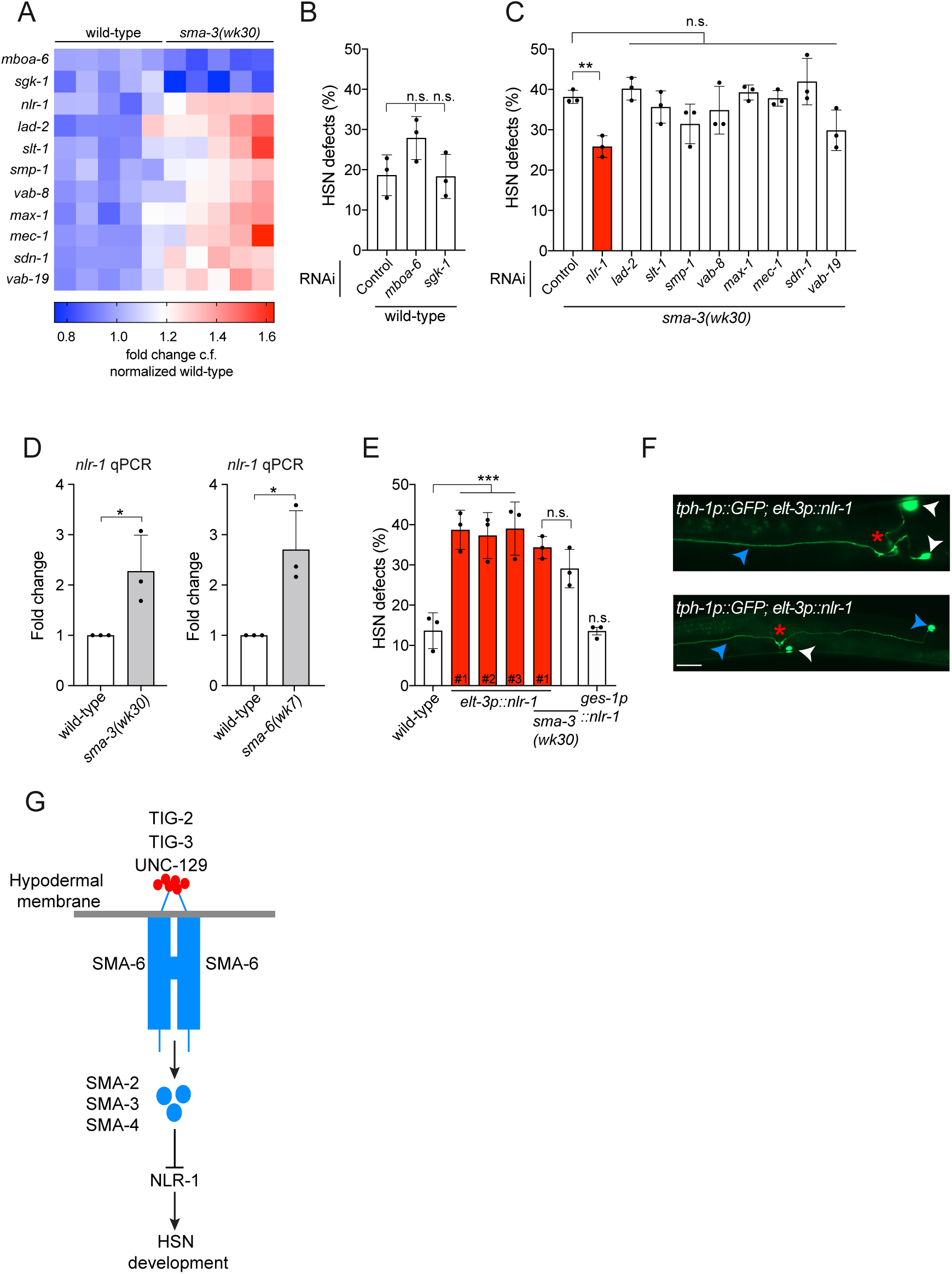
SMA-3 regulation of the NLR-1/Neurexin-like receptor controls HSN development. (A) Heat map of whole-animal transcriptional profiling from *sma-3(wk30)* L2 larvae compared to wild-type. Gene names of candidate HSN regulators on the left. Each column represents individual total RNA samples. Downregulated (blue), upregulated (red) and unchanged (white). (B) Quantification of HSN developmental defects in wild-type animals following RNAi knockdown of genes downregulated in the *sma-3(wk30)* mutant. n>100; n.s. not significant (One-way ANOVA with Tukey’s correction). Error bars represent mean ± SEM. (C) Quantification of HSN developmental defects in *sma-3(wk30)* animals following RNAi knockdown of genes upregulated in the *sma-3(wk30)* mutant. n>100; **P<0.01, n.s. not significant (One-way ANOVA with Tukey’s correction). Error bars represent mean ± SEM. (D) Relative *nlr-1* mRNA levels in *sma-3(wk30)* (left) and *sma-6(wk7)* (right) mutant L2 larvae normalized to values for wild-type worms. Three biological replicates were compared (*cdc-42* reference gene was used). *P<0.05 (t test). Error bars represent mean ± SEM. (E) Quantification of HSN developmental defects in animals overexpressing *nlr-1* in the hypodermis (*elt-3* promoter) or intestine (*ges-1* promoter). *nlr-1* overexpression in the hypodermis but not the intestine causes HSN defects. HSN defects of *sma-3(wk30)* animals are not enhanced by *nlr-1* overexpression in the hypodermis. n>100; ***P<0.001, n.s. not significant (One-way ANOVA with Tukey’s correction). Error bars represent mean ± SEM. # refers to independent transgenic lines (line #1 in the *sma-3(wk30)* background is same line used in wild-type). (F) HSN developmental defects caused by overexpression of *nlr-1* in the hypodermis are phenotypically similar to those observed with loss of *sma-6* and *sma-3* (see Fig. 1B and Table S1). Vulval position is marked with a red asterisk, wild-type positioned cell bodies with white arrowheads, and misguided cell bodies and axons with blue arrowheads. Ventral view, anterior to the left. Scale bar: 20 μm. (G) The TGF-β ligands TIG-2, TIG-3 and UNC-129 regulate the TGF-β type I receptor SMA-6 to control HSN guidance non-cell-autonomously from the hypodermis. Hypodermal TGF-β signaling through the SMAD-2/3/4 transcriptional regulators limits expression of the Neurexin-like cell adhesion molecule NLR-1 to enable faithful HSN development.

NLR-1 is member of Caspr subfamily of neurexin-like proteins (*26*). This family of transmembrane cell adhesion molecules mediate neuron-neuron interactions and functionally organize synapses (*27, 28*). NLR-1 contains extracellular Laminin G, epidermal growth factor and fibrinogen-related domains, suggesting roles in cell adhesion and cell-cell signaling (*26*). A *C. elegans nlr-1* deletion mutant (*tm2050)* causes embryonic lethality (*C. elegans* Gene Knockout Consortium), demonstrating the essential requirement for this protein (*29*). Our RNAi knockdown experiments show that reducing *nlr-1* expression suppresses the HSN defects caused by loss of *sma-3*, suggesting that SMA-6 signaling limits *nlr-1* expression (Fig. 4C). We tested this hypothesis by comparing *nlr-1* mRNA abundance in *sma-3(wk30)* and *sma-6(wk7)* mutants compared to wild-type animals. Using quantitative polymerase chain reaction analysis, we found that *nlr-1* expression is increased by approximately two-fold in the absence of either *sma-3* (as revealed by RNA-seq) or *sma-6* (Fig. 3D). These data show that SMA-6 signaling limits *nlr-1* expression.

Due to the hypodermal-specific function of SMA-6 in non-cell-autonomously regulating HSN development (Fig. 1C), we predicted that overexpression of *nlr-1* in the hypodermis, and not in another tissue, would cause HSN defects in wild-type animals. We found that *nlr-1* overexpression in the hypodermis (*elt-3* promoter), but not the intestine (*ges-1* promoter), caused HSN defects similar to those observed in the *sma-3(wk30)* mutant both quantitatively and qualitatively (Fig. 4E-F and Table S1). Importantly, hypodermal overexpression of *nlr-1* did not further exacerbate *sma-3(wk30)* HSN defects (Fig. 4E). These data reveal that SMA-6 signaling in the hypodermis prevents expression of the neurexin-like protein NLR-1 to optimize the extracellular landscape, and enable correct HSN development (Fig. 4G).

We have shown here that the SMA-6 TGF-β type I receptor controls a developmental decision independently of DAF-4 - the sole TGF-β type II receptor in *C. elegans*. To our knowledge, this is the first report of such a function for a TGF-β type I receptor. A previous report did reveal, however, that the other *C. elegans* TGF-β type I receptor, DAF-1, acts partially in parallel with DAF-4 to control dauer formation (*20*). Our data show that SMA-6 acts in the hypodermis to control HSN guidance. DAF-4 is also expressed in this tissue (*20*), suggesting that SMA-6 regulates HSN guidance from distinct membrane domains or cellular subpopulations within the hypodermis. Intriguingly, we also show that three, orphan, TGF-β ligands (TIG-2, TIG-3 and UNC-129) physically interact with SMA-6, and act in the same genetic pathway with each other and SMA-6 to control HSN development. TIG-2 functions from the nervous system and TIG-3/UNC-129 from BWM in this regard. This posits that TIG-2 homodimers and TIG-3/UNC-129 homo- or heterodimers form *in vivo* and interact with SMA-6, potentially in a complex with an unknown receptor(s). A previous study found a similar non-redundant requirement for three Activin ligands (TGF-β superfamily members) in the *Drosophila* retina (*30*). Further, cell-based assays showed that promiscuous interactions between TGF-β superfamily ligands and their receptors enable cells to perceive information encoded by distinct ligand combinations (*31*). Thus, discrete intracellular responses are dependent on ligand composition and concentration. We found here that the intracellular responses downstream of SMA-6 are coordinated by the SMA-2/3/4 SMAD complex, that represses expression of NLR-1, a neurexin-like cell adhesion receptor. Limiting NLR-1 expression is important as its inappropriate expression in the hypodermis causes defective neuronal migration and axon guidance, presumably due to aberrant extracellular adhesion and/or signalling. Together, our results imply that correct brain development requires tight regulation of Neurexin expression to ensure an optimal adhesive and architectural environment for navigating neurons.

## Acknowledgments

We thank members of Pocock laboratory and Brent Neumann for comments on the manuscript. Some strains used in this study were provided by the *Caenorhabditis* Genetics Center, which is funded by NIH Office of Research Infrastructure Programs (P40 OD010440), and by Shohei Mitani at the National Bioresource Project (Japan). We also thank Rick Padgett for the *elt-3p::sma-6::gfp* plasmid, Jun Liu for the Myc-SMA-6 plasmid, Yuichi Iino for the *rimb-1* promoter and Yun Zhang for the *sma-6(wk7); ges-1p::sma-6* and *sma-6(wk7); sma-6p::sma-6* rescue strains. The authors also acknowledge use of the services and facilities of Micromon Genomics at Monash University.

## Author contributions

O.B., M.E.P., T.S., A.H., G.S., M.H. and R.P. performed experiments. S.A. analyzed the RNA sequencing data. R.P. supervised the research and wrote the manuscript.;

## Data and materials availability

RNA-seq data has been deposited into the NCBI Gene Expression Omnibus (GEO) under accession number GSE151035. All data and materials used in the analysis must be available in some form to any researcher for purposes of reproducing or extending the analysis.

